# Anatomy of unfolding: The site-specific fold stability of Yfh1 measured by 2D NMR

**DOI:** 10.1101/2021.03.27.437329

**Authors:** Rita Puglisi, Gogulan Karunanithy, D. Flemming Hansen, Annalisa Pastore, Piero Andrea Temussi

## Abstract

Most techniques allow detection of protein unfolding either by following the behaviour of single reporters or as an averaged all-or-none process. We recently added 2D NMR spectroscopy to the well-established techniques able to obtain information on the process of unfolding using resonances of residues in the hydrophobic core of a protein. Here, we questioned whether an analysis of the individual stability curves from each resonance could provide additional site-specific information. We used the Yfh1 protein that has the unique feature to undergo both cold and heat denaturation at temperatures above water freezing at low ionic strength. We show that stability curves inconsistent with the average NMR curve from hydrophobic core residues mainly comprise exposed outliers that do nevertheless provide precious information. By monitoring both cold and heat denaturation of individual residues we gain knowledge on the process of cold denaturation and convincingly demonstrate that the two unfolding processes are intrinsically different.

## Introduction

We are all accustomed to the concept that proteins unfold when temperature is increased. Less well known is that all proteins unfold in principle also at low temperatures as demonstrated by P. Privalov on purely thermodynamics grounds (Privalov, 1990). According to this theory, the driving force of heat denaturation is the increase of conformational entropy with temperature. This automatically involves the hydrophobic core and disfavours less ordered parts of the architecture. On the contrary, while the mechanism of cold denaturation is still debated, the current hypothesis is that this transition occurs when entropy decreases. In this case, the driving force of unfolding would be driven by the sudden solvation of the hydrophobic residues of the core (Privalov, 1990).

The reason why cold denaturation is much less understood than the heat transition is that most proteins undergo cold denaturation at temperatures below the water freezing point. This is unfortunate because observation of both unfolding temperatures is in principle very valuable as it allows calculation of reliable stability curves of the protein and of the whole set of thermodynamic parameters.

We have identified a protein, Yfh1, that, as a full-length natural protein, undergoes cold and heat denaturation at detectable temperatures when in the absence of salt (Pastore et al., 2007). We have extensively exploited these properties to gain new insights both on the denatured states of Yfh1 (Adrover et al., 2012) and on the factors that may influence its stability (Martin et al., 2008). The value of Yfh1 as a tool to investigate the unfolding process is evidenced not only by our subsequent work (Sanfelice et al., 2013; Pastore and Temussi, 2017; Sanfelice et al., 2014; Alfano et al., 2017) but also by papers from other laboratories (Espinosa et al., 2016; Chatterjee et al., 2014; Bonetti et al., 2014; Aznauryan et al., 2013).

In our studies, we noticed that most techniques employed to monitor protein stability are however not “regiospecific”, as they yield a global result, *i*.*e*. an estimate of the stability of the whole protein architecture, observable through the global evolution of secondary structure elements upon an environmental insult. This is because we postulate an all-or-none cooperative process in which the protein collapses altogether from a folded to an unfolded state. When monitoring unfolding of a protein by CD spectroscopy, for instance, we observe intensity changes related to the disruption of alpha helices and/or beta sheets under the influence of physical or chemical agents.

It would instead be interesting to gauge the response of selected regions of the protein at the single residue level to gain new insights into the mechanisms of unfolding of selected parts of the protein structure. A technique ideally suited for this purpose is 2D ^15^N HSQC spectroscopy since it provides a direct fingerprint of the protein through mapping each of the amide protons. Volume variations of the NMR resonances may reflect changes affecting single atoms of each residue and indirectly report on how they are individually affected by the unfolding process. We recently showed, using Yfh1 as a suitable model, that it is possible to use 2D NMR to measure protein stability and get thermodynamic parameters comparable to those obtained by CD (Puglisi et al., 2020). We showed that this is possible provided that the residues chosen are those buried in the hydrophobic core, thus experiencing the unfolding process directly. To reliably select these residues, we introduced a parameter RAD which was defined as the combination of the depth of an amide group from the protein surface and the relative accessibility at the atom level (Puglisi et al., 2020). We demonstrated that, by excluding most of the exposed residues (RAD values for the amide nitrogens ≥ 0.5) and averaging over resonances from residues with RAD values lower than 0.1, we can obtain thermodynamics parameters indistinguishable, within experimental error, from those obtained by CD or 1D NMR (Puglisi et al., 2020).

Using the approach previously developed (Puglisi et al, 2020), we systematically analysed in the current work the heat and cold denaturation of Yfh1 at residue detail but we reversed the perspective and wondered what information, if any, would be carried by residues far from the hydrophobic core and how they reflect the process of unfolding. This subject has increasingly attracted attention: as put in the words of a recent study by Grassein et al. (2020): “For most of the proteins, this global heat-induced denaturation curve can be formally described by a simple two-state (folded/unfolded) statistical model. Agreement with a two-state model does not imply, however, that the macromolecule does not unfold through a number of intermediate states…. Hence, the global denaturation curve hides the heterogeneity of protein unfolding. …Local nativeness is not uniquely defined and is probe dependent.” Understanding how individual residues report on protein unfolding is also relevant in view of an increasing number of studies on protein stability based on the intensity variations of the resonance of a single residue upon unfolding (Danielsson et al., 2015; Smith et al., 2016; Guseman et al., 2018). The excellent agreement between NMR and CD thermodynamic parameters using 2D NMR (Puglisi et al., 2020) put us in the position to examine the output of single residues critically and follow the process of unfolding at an atomic level.

Using once again Yfh1, we show here that it is possible to sort out which individual single residues yield stability curves consistent with the global unfolding process and that we can obtain valuable information on the process of unfolding from residues that diverge from the average behaviour: whereas some of the residues signal a single folding/unfolding event, we find that others report on more complex thermodynamic events. Our data directly demonstrate that the cold and heat denaturation processes have distinctly different mechanisms and provide site-specific information on solvent interactions supporting Privalov’s interpretation of cold denaturation (Privalov, 1990). Our results also clearly demonstrate the considerable advantages of NMR over other approaches, such as in CD or fluorescence, that probe only bulk transitions or individual residues.

## Results

### Data collection and preliminary considerations

To study the unfolding of Yfh1, we collected ^15^N HSQC spectra of Yfh1 at different temperatures and extracted the volumes of individual residues as a function of temperature (**Figure S1 of Suppl. Mat**.). This could be confidently done for 68 (out of the expected 109) well resolved resonances. The behaviour of ^15^N HSQC spectra of Yfh1 as a function of temperature was not uniform: some peaks could be observed nearly at all temperatures in the range 273-323 K, others disappeared at temperatures intermediate between room temperature and the two unfolding temperatures, *i*.*e*. lower than 323 K or higher than 273 K (**Figures S2 and S3 of Suppl. Mat**.). This behaviour can of course be ascribed to the exchange regime (intermediate) between folded and unfolded conformations of these residues and told us that they are not an integral part of the architecture of the folded form. The possibility that the intensity changes in the HSQCs at low temperature could be solely due to exchange broadening and not to unfolding can however be excluded by the practically perfect agreement between the curves obtained by CD and by NMR (both 1D (Pastore et al., 2007) and 2D (Puglisi et al., 2020)). Cold denaturation of Yfh1 has also been independently confirmed by five independent techniques (Espinosa et al., 2016; Chatterjee et al., 2014; Bonetti et al., 2014; Aznauryan et al., 2013).

### Extraction of the thermodynamics parameters

We could then extract the thermodynamic parameters of the unfolding process for the selected resonance assuming that some conditions are met (Privalov, 1990; Martin et al., 2008). We first assumed that unfolding transitions are, at a first approximation, two-state processes from folded (F) to unfolded (U) states. We then postulated that the difference of the heat capacity of the two forms (ΔC_p_) does not depend on temperature. This assumption is considered reasonable when the heat capacities of the native and denatured states change in parallel with temperature variations (Privalov, 1990). When these two conditions are reasonably met, the populations of the two states at temperature T, f_F_(T) and f_U_(T), are a function of the Gibbs free energy of unfolding, ΔG°(T) (see Methods and Martin et al., 2008). The plot of the free energy of unfolding as a function of temperature provides what is called the stability curve of a protein (Becktel and Schellman, 1987). From this equation the main thermodynamic parameters, i.e. heat melting temperature (T_m_), enthalpy difference at the melting point (ΔH_m_) and the heat capacity difference at constant pressure (ΔC_p_), can be determined using a non-linear fit (damped least-squares method, also known as the Levenberg-Marquardt algorithm) (Levenberg, 1944; Marquardt, 1963). Other parameters, *e*.*g*. the low temperature unfolding (T_c_), can be read from the stability curve. When the original assumptions are significantly wrong, fitting results in unrealistic numbers. In our case, the volumes were transformed into relative populations of folded Yfh1 assuming that, as measured by CD and confirmed in other studies on Yfh1 (Pastore et al., 2007; Martin et al., 2008; Sanfelice et al., 2014; Alfano et al., 2017), unfolded forms are in equilibrium with, on average, a 70% population of folded Yfh1 at room temperature. The concurrent presence of an equilibrium between folded and unfolded species of Yfh1 at low ionic strength was proven by the co-existence of minor extra peaks which disappear as soon as physiologic concentrations of salt are added (Vilanova et al., 2014).

### Identification of residues consistent with or outliers from the global behaviour

We correlated each amide resonance to the corresponding value of RAD, the parameter introduced in Puglisi et al. (2020), to pinpoint residues close to the hydrophobic core (**Table 1**). Of the 68 residues selected, 39 had RAD <0.5, 37 with RAD <0.4, 33 with RAD <0.3, 24 with RAD <0.2 and 11 RAD < 0.1 (**Table 1**). The residues with RAD<0.1 (henceforth called RAD_0.1) were used to calculate the average. Comparison of the stability curves of the non-overlapping amide resonances with this average showed that several residues with quite different RAD values yield stability curves drastically different from the average (**Figures 1a**). We next tried to classify the individual stability curves into those that matched well the average RAD_1 curve (‘well-behaved’) and those that did not (‘ill-behaved’). The curves for residues in the hydrophobic core were in good agreement with the average curve (**Figures 1b**).

**Table 1.** Thermodynamic parameters of the detectable residues. The average stability curve was obtained selecting the residues with RAD_0.1 indicated in the table in bold face (Puglisi et al., 2020).

| | $\Delta H$ (Kcal/mol) | $\Delta S$ (Kcal/mol) | $\Delta C_p$ (Kcal/molK) | $T_m$ (K) | $T_c$ (K) | RAD |
| --- | --- | --- | --- | --- | --- | --- |
| 61 Val | 19,9 | 0,067 | 1,58 | 298,4 | 273,9 | 48,20 |
| 63 Gln | 21,1 | 0,072 | 2,24 | 294,0 | 275,6 | 3,64 |
| 64 Glu | 27,7 | 0,093 | 3,30 | 298,0 | 281,5 | 2,52 |
| 65 Val | 24,3 | 0,081 | 2,71 | 299,7 | 282,2 | 0,31 |
| 68 Leu | 21,0 | 0,071 | 3,02 | 296,5 | 282,8 | 0,75 |
| 70 Leu | 28,5 | 0,096 | 3,73 | 298,2 | 283,1 | 2,35 |
| 71 Glu | 29,1 | 0,097 | 3,59 | 299,8 | 283,9 | 7,13 |
| 72 Lys | 33,1 | 0,111 | 3,73 | 299,6 | 282,2 | 6,24 |
| 75 Glu | 24,3 | 0,081 | 2,75 | 300,0 | 282,7 | 2,07 |
| 76 Glu | 17,4 | 0,058 | 1,71 | 300,6 | 280,8 | 0,91 |
| 78 Asp | 29,0 | 0,096 | 3,18 | 301,1 | 283,2 | 0,17 |
| 83 His | 22,8 | 0,076 | 2,21 | 298,4 | 278,2 | 0,21 |
| 86 Asp | 25,4 | 0,084 | 2,62 | 301,0 | 282,0 | 0,27 |
| 87 Ser | 22,8 | 0,076 | 2,42 | 299,1 | 280,6 | 0,34 |
| **88 Leu** | **20,2** | **0,067** | **1,75** | **300,6** | **278,1** | **0,04** |
| 89 Glu | 22,1 | 0,074 | 2,14 | 299,7 | 279,5 | 0,20 |
| 90 Glu | 26,1 | 0,087 | 2,50 | 300,9 | 280,5 | 0,52 |
| 91 Leu | 34,1 | 0,114 | 4,19 | 300,3 | 284,3 | 0,16 |
| 92 Ser | 22,6 | 0,075 | 1,97 | 299,7 | 277,3 | 0,15 |
| 93 Glu | 19,5 | 0,065 | 1,97 | 300,1 | 280,7 | 0,61 |
| 94 Ala | 23,6 | 0,079 | 2,55 | 299,6 | 281,5 | 4,10 |
| 95 His | 18,6 | 0,062 | 1,31 | 300,7 | 273,2 | 0,28 |
| 97 Asp | 23,3 | 0,078 | 2,52 | 299,0 | 280,9 | 0,95 |
| 98 Cys | 22,6 | 0,076 | 2,59 | 297,7 | 280,6 | 0,26 |
| 99 Ile | 17,6 | 0,059 | 1,75 | 297,5 | 277,8 | 0,11 |
| 101 Asp | 22,3 | 0,074 | 2,42 | 300,2 | 282,1 | 1,08 |
| 104 Leu | 29,5 | 0,098 | 3,12 | 300,9 | 282,4 | 0,78 |
| 105 Ser | 23,7 | 0,079 | 2,51 | 299,7 | 281,2 | 1,18 |
| 107 Gly | 19,3 | 0,065 | 2,44 | 296,6 | 281,1 | 3,58 |
| 108 Val | 22,3 | 0,075 | 2,03 | 299,0 | 277,5 | 0,50 |
| 109 Met | 26,6 | 0,089 | 3,25 | 299,4 | 283,3 | 0,63 |
| 110 Thr | 33,2 | 0,111 | 3,79 | 300,2 | 283,0 | 0,23 |
| 112 Glu | 28,5 | 0,094 | 2,88 | 301,9 | 282,5 | 0,41 |
| 113 Ile | 17,4 | 0,059 | 2,11 | 297,6 | 281,3 | 0,12 |
| 115 Ala | 15,4 | 0,052 | 2,77 | 295,8 | 284,9 | 2,48 |
| 116 Phe | 14,8 | 0,049 | 1,73 | 298,0 | 281,3 | 0,62 |
| 117 Gly | 27,6 | 0,093 | 3,48 | 298,2 | 282,6 | 0,98 |
| 119 Tyr | 24,9 | 0,083 | 2,72 | 300,3 | 282,3 | 0,22 |
| 120 Val | 15,7 | 0,053 | 1,68 | 297,3 | 279,0 | 0,33 |
| 127 Asn | 23,0 | 0,077 | 2,46 | 297,0 | 278,7 | 5,81 |
| 128 Lys | 15,5 | 0,052 | 1,35 | 300,4 | 277,9 | 0,66 |
| 129 Gln | 14,3 | 0,048 | 1,80 | 296,8 | 281,2 | 0,20 |
| **130 Ile** | **27,9** | **0,093** | **3,19** | **300,9** | **283,8** | **0,02** |
| **131 Trp** | **21,7** | **0,073** | **2,54** | **298,6** | **281,8** | **0,04** |
| **132 Leu** | **26,4** | **0,088** | **3,34** | **300,7** | **285,1** | **0,02** |
| 133 Ala | 24,7 | 0,082 | 2,52 | 299,9 | 280,7 | 0,19 |
| 134 Ser | 20,3 | 0,068 | 2,07 | 299,7 | 280,5 | 0,13 |
| 136 Leu | 13,2 | 0,044 | 1,27 | 299,3 | 278,9 | 0,25 |
| 140 Asn | 25,4 | 0,085 | 2,80 | 299,4 | 281,6 | 0,17 |
| **142 Phe** | **20,9** | **0,070** | **3,06** | **299,3** | **285,8** | **0,03** |
| 143 Asp | 21,9 | 0,073 | 2,50 | 300,2 | 283,1 | 0,13 |
| 146 Asn | 23,6 | 0,080 | 3,61 | 295,0 | 282,1 | 2,00 |
| 147 Gly | 25,2 | 0,085 | 2,37 | 297,8 | 277,0 | 4,80 |
| 148 Glu | 21,6 | 0,072 | 2,69 | 298,8 | 283,0 | 1,40 |
| **150 Val** | **22,9** | **0,076** | **2,20** | **300,7** | **280,4** | **0,03** |
| **151 Ser** | **22,7** | **0,076** | **2,39** | **299,9** | **281,3** | **0,05** |
| 152 Leu | 32,2 | 0,107 | 3,87 | 300,0 | 283,7 | 0,16 |
| 154 Asn | 21,9 | 0,074 | 2,40 | 295,1 | 277,2 | 1,14 |
| **158 Leu** | **29,1** | **0,096** | **3,11** | **301,9** | **283,6** | **0,03** |
| **159 Thr** | **29,8** | **0,099** | **3,38** | **300,0** | **282,8** | **0,09** |
| 160 Asp | 23,6 | 0,078 | 2,55 | 301,2 | 283,1 | 0,28 |
| **161 Ile** | **15,4** | **0,051** | **1,66** | **300,1** | **281,9** | **0,09** |
| 163 Thr | 15,2 | 0,051 | 1,44 | 300,9 | 280,2 | 0,15 |
| **166 Val** | **27,3** | **0,091** | **2,81** | **300,8** | **281,8** | **0,06** |
| 168 Lys | 17,3 | 0,058 | 1,93 | 299,9 | 282,4 | 0,16 |
| 170 Ile | 22,6 | 0,075 | 2,11 | 301,3 | 280,3 | 0,31 |
| 172 Lys | 28,1 | 0,095 | 3,84 | 294,1 | 279,7 | 1,5 |
| 174 Gln | 20,6 | 0,069 | 2,2 | 297,5 | 279,2 |  |
| 131 Trp sc | 27,1 | 0,091 | 3,21 | 299,4 | 282,8 |  |

**Figure 1.**
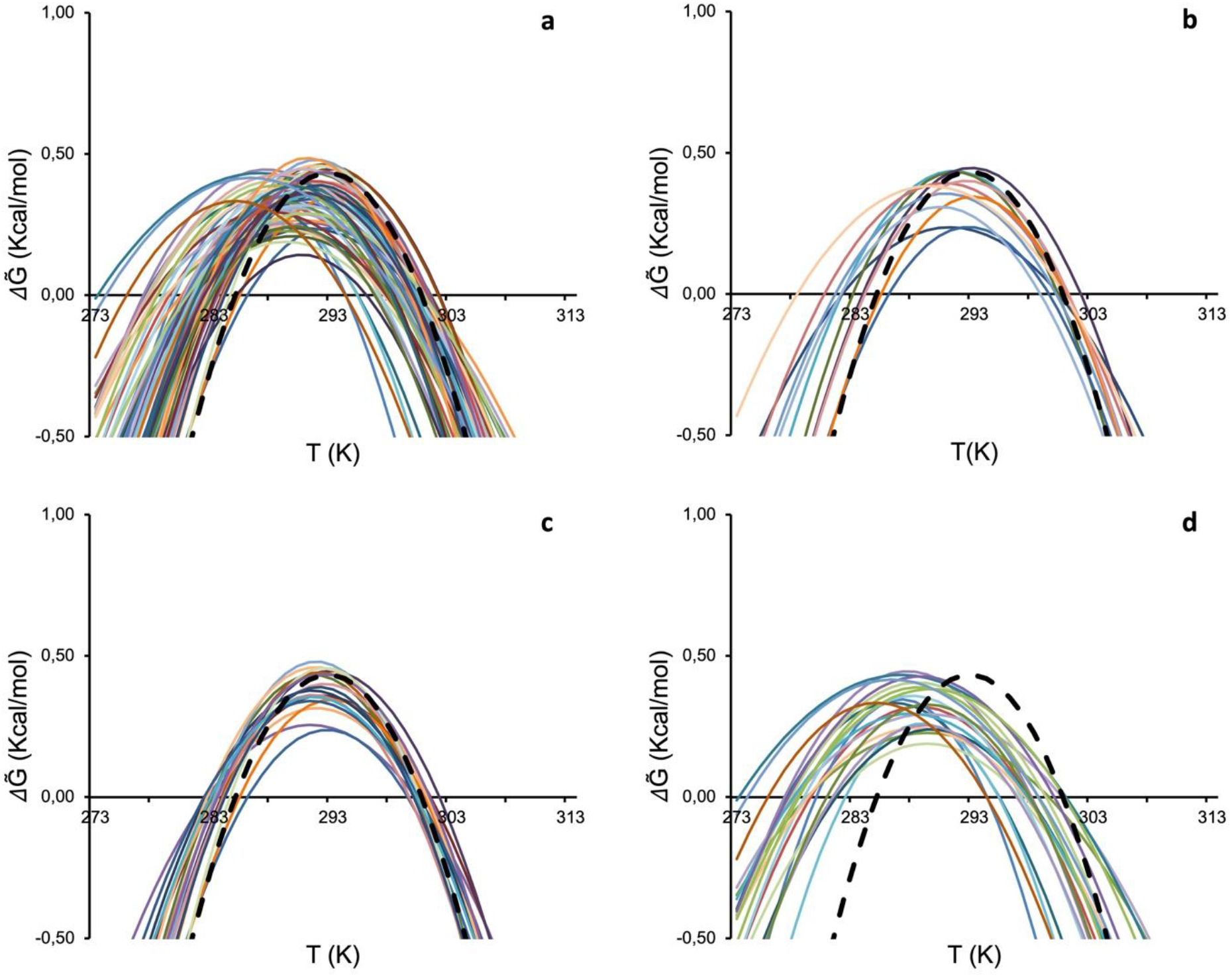
Comparison of single residue stability curves with the global RAD_0.1 best curve (dashed black). **a**) Stability curves of all observable isolated residues. **b**) Stability curves of residues with a RAD<0.1. **c**) Stability curves of single residues for which the difference in the unfolding temperatures with respect to values of the reference curve (ΔT_m_ and ΔT_c_) is on average below 1.5 °C **d**) Stability curves of single residues for which the difference in the unfolding temperatures with respect to values of the average curve (ΔTm and ΔTc) is on average above 3 K. For simplicity, colour coding is not the same in the different panels.

However, we could not find in general a clear-cut criterion to decide when the curves were not consistent with the average. We arbitrarily chose to set a cut-off at values of the unfolding temperatures (T_m_ and T_c_) that differed, on average, less than 1.5 K from those corresponding to the average (RAD_0.1). These ΔT_m_ and ΔT_c_ differences are smaller than the variability that we had observed among different preparations and measurements of the same protein (Pastore et al., 2007; Martin et al., 2008; Sanfelice et al., 2014; Sanfelice et al., 2015; Alfano et al., 2017; Puglisi et al., 2020). The residues selected according to this criterion are E71, E75, D78, L91, D101, L104, M109, T110, Y119, I130, L132, F142, D143, L152, L158, T159, D160 and K168 (**Figure 1c**). Most of the amide groups of the well-behaved residues are spread among well-structured secondary elements, but a few are in less ordered regions (**Figure 2a)**. By the same token, we selected as ‘ill-behaved’ residues those whose T_m_ and T_c_ values differed from the average curve, on average, more than 3 K with respect to the best curve RAD_0.1. Twenty-one residues (V61, Q63, H83, L88, S92, H95, C98, I99, G107, V108, I113, V120, N127, K128, Q129, L136, N146, G147, N154, K172, Q174) belong to this sub-set. Except for a few outliers, they are all in less structured regions (**Figure 2b)**. Amongst these residues, V61, Q63, H95 which are positioned in flexible regions (either in the N-terminal tail or in a loop), are those with the largest shift of T_c_. This behaviour is, however, not a general rule as some of the best-behaved residues reported in **Figure 1c** are not in regular secondary structure elements confirming the complexity of the system under study.

**Figure 2.**
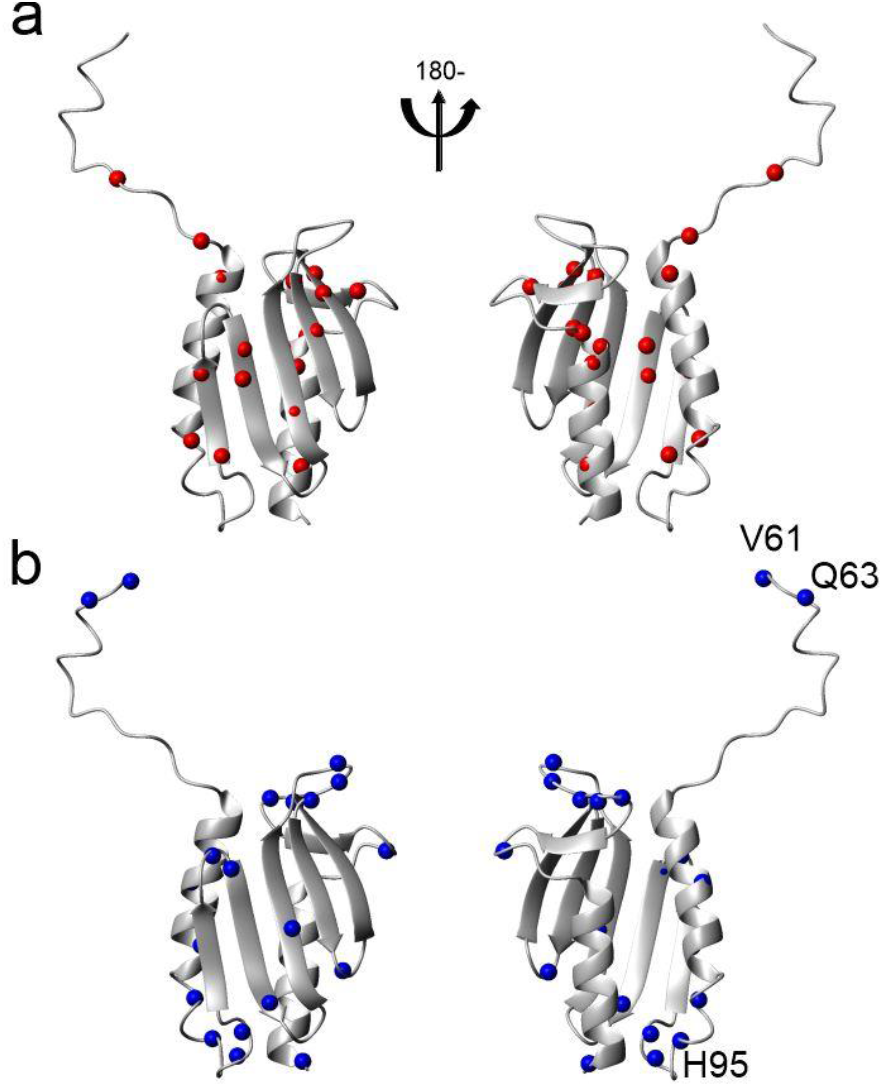
Distribution of residues on the structure of Yfh1 (pdb id 2fql). a) Distribution of the nitrogen atoms of residues for which the difference in the unfolding temperatures with respect to values of the RAD_0.1 curve (ΔTm and ΔTc) is on average below 1.5 K. b) Distribution of the N atoms of residues for which the difference in the unfolding temperatures with respect to values of the average curve (ΔTm and ΔTc) is on average above 3 K. Indicated explicitly are the three residues whose stability curve is most shifted to lower temperatures with respect to the average RAD_0.1. The structure pairs are rotated by 180 degrees around the y axis.

The stability curves of the residues that differ from the average (**Figure 1d**) have an important peculiarity: most stability curves show a moderate decrease of T_m_ (ΔT_m_<0) and a large decrease of T_c_ (ΔT_c_<<0) from the average. This finding would imply that the corresponding transition temperatures for the heat and cold unfolding point to a decreased stability for heat denaturation but an increased stability for cold denaturation.

### Evaluating the contribution of errors

To make sure that the effect is beyond experimental errors, we reasoned that three phenomena could potentially lead to erroneous populations, f_F_(T) and f_U_(T), and thus stability curves: 1) the folding exchange dynamics leading to a time-dependent fluctuation of the ^1^H chemical shift and loss of intensity during the INEPTs of the ^15^N-HSQC, 2) differential intrinsic relaxation rates in the folded and unfolded states, and 3) exchange of the detected amide protons with the bulk solvent. We thus performed simulations to evaluate how much these phenomena could influence the resulting curves (for a more detailed discussion see **Suppl. Mat**.). We found that, although the three contributions affect the derived populations, the stability curves that are naïvely calculated from the intensities observed in the NMR spectra as 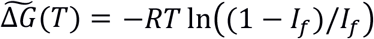, where *I*_f_ is the peak intensity of the folded species, recapitulate the general features of the expected stability curve, Δ*G(T)*. Of particular interest is that the temperature of maximum stability T_S_ (so called because it corresponds to zero entropy of the stability curve), is well reproduced despite the deviations observed for the other parameters (**Figure S4 of Suppl. Mat**.).

Our observations are thus beyond experimental error and indicate that the mechanisms of the two unfolding processes, at high and low temperatures, are intrinsically different in agreement with Privalov’s theory (Privalov, 1990).

### A possible classification of the outliers

The negative values of ΔT_m_ and ΔT_c_ observed for some residues (**Figure 1d**) imply that also the temperature of maximum stability T_S_ for these residues is lower than that observed for the best average RAD_0.1. A shift of T_S_ towards higher temperature values, when studying several cases of thermophilic proteins, was attributed by Razvi & Scholtz (2006) to a decrease in the entropy difference in unfolding. Obviously, a *decrease* of T_m_ or T_c_ caused by shifting the Ts to lower temperatures is connected to an increase in the entropy difference. This interpretation is based on the classification by Nojima et al. (1977) of the main mechanisms of changing the thermal resistance, that is the resistance of heat to cross a material, of a protein. According to the rough classification of Nojima et al. (1977), altered thermostability can be achieved thermodynamically in three extreme cases (**Figure 3**). Real situations might of course contain mixtures of the three possibilities.

**Figure 3.**
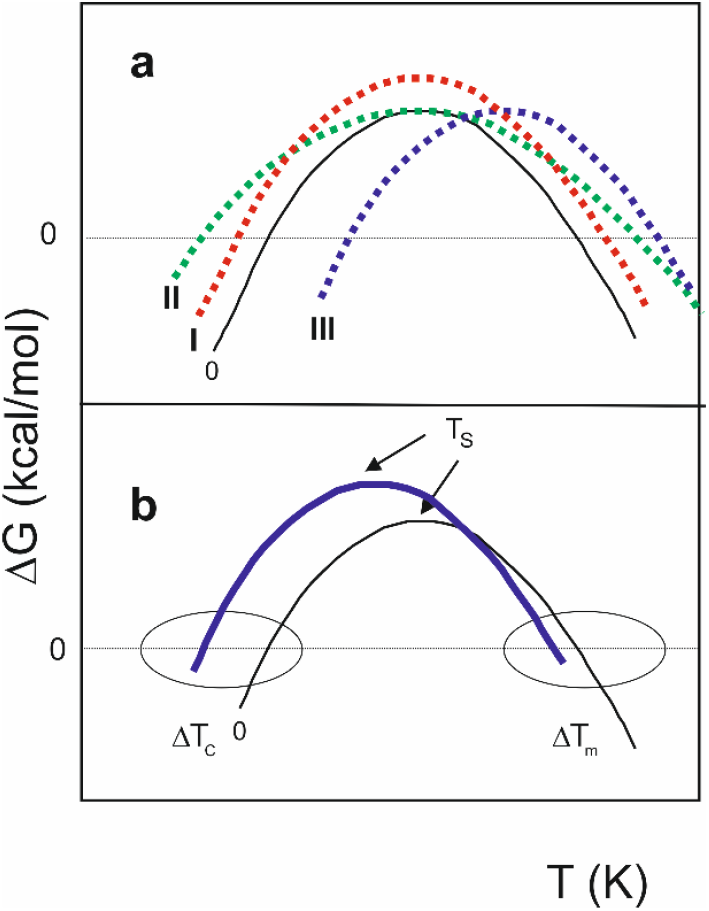
Mechanisms that influence stability curves of a protein (adapted from Nojima et al, 1977). **a**) Dependence of the difference of free energy between unfolded and folded states (**Δ**G) of a hypothetical protein vs temperature (T) (curve 0). Mechanism I illustrates the effect of increasing **Δ**H_S_ (curve I). Mechanism II shows the effect of reducing **Δ**C_p_ (curve II). Mechanism III shows the shift of the whole stability curve towards higher temperatures caused by decreasing **Δ**S_m_ (curve III). **b**) A combination of the three mechanisms. The solid blue curve, with a prevalent low shift of T_S_, corresponds qualitatively to the cases of Yfh1 reported in **Figure 1d**.

According to mechanism I, when ΔH_S_ (the change in enthalpy measured at T_S_) increases, the stability curve retains the same shape, but with greater ΔG values at all temperatures. With mechanism II, a decreased ΔC_p_ leads to a broadened stability curve retaining the same maximum, because the curvature of the stability curve is given by 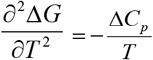 (Becktel, & Schellman, 1987). According to mechanism III, the entire curve can shift towards higher or lower temperatures. It is possible to show (Privalov, 1990) that:

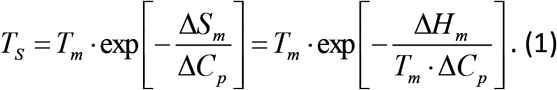

Increasing the difference in entropy between the folded and unfolded states (ΔS_m_) can shift values of T_S_ towards lower temperatures. Most of the curves in **Figure 1d** do not correspond to a single mechanism, but to a combination of them (**Figure 3b**). Nevertheless, all curves are shifted towards lower values of T_S_ and larger low-temperature differences correlate well with less ordered regions of the structure. It is thus not surprising to find this behaviour for residues at the N- and C-termini (Q63 and K172) or in connecting loops (G107, N127, N146 and N154) which are bound to be flexible (Halle, 2002). More surprising is, however, to find amongst these residues also V120 which is right in the middle of the beta sheet. While we have not a definite explanation for this observation at the moment, it could indicate a local frustration point in this region.

### Exploring the correlation between stability and secondary structure elements

We have previously shown that, in addition to the criteria of depth and exposition, an alternative selection of residues over which average populations might be based on elements of regular secondary structure (Puglisi et al., 2020). It is now possible to analyse the behaviour of each secondary structure element. Of the 68 residues selected, 35 were in secondary structure elements (15 in alpha helices, 20 in beta sheets). The largest number of residues of secondary structure traits whose resonance is accessible belongs to the two helices (**Figure 4**).

**Figure 4.**
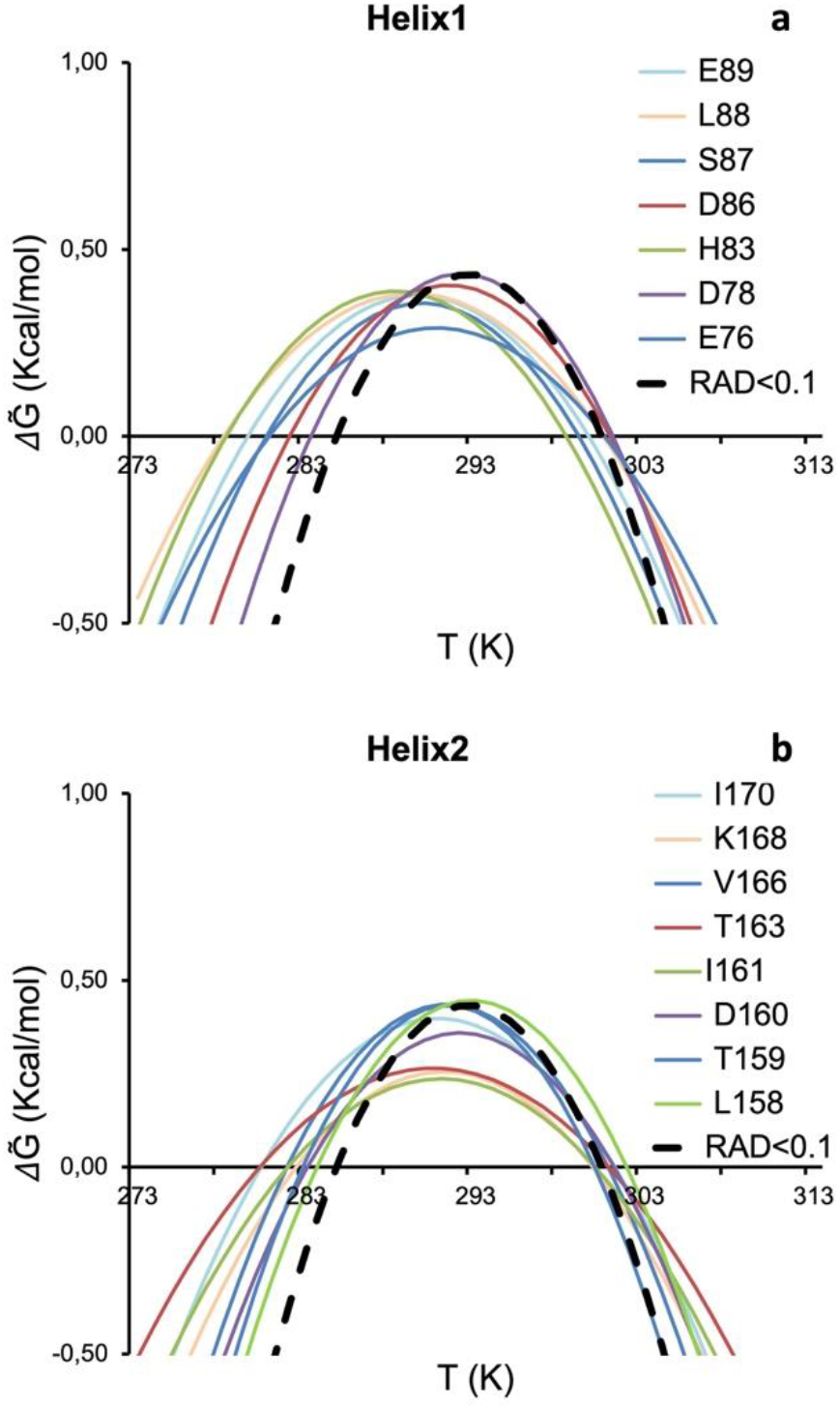
Stability curves of residues belonging to secondary structure elements. **a**) Helix 1. **b**) Helix 2. Residues are labelled with single letter code. The average stability curve is shown as black dashed line.

Several resonances have stability curves far from the reference one (dashed black curve of RAD_0.1). These are those of His83 and Leu88 for helix 1 (**Figure 4a**). All the others are in fair agreement with the average curve. The best-behaved residue (Asp78) is located at the end of the helix with its amide groups in the buried side of the helix. For helix 2, the worst agreement is found for Thr163 and Ile170, whereas the best agreement is for Leu158, Thr159, Asp160 and Lys168 (**Figure 4b**). This implies that residues of helix 2 with a good agreement are distributed over the whole secondary structure element. Some residues of helix 2 have also lower stability curves which indicate a lower ΔH.

The number of residues belonging to beta strands for which it was possible to extract stability curves is more limited (**Figure 5)**.

**Figure 5.**
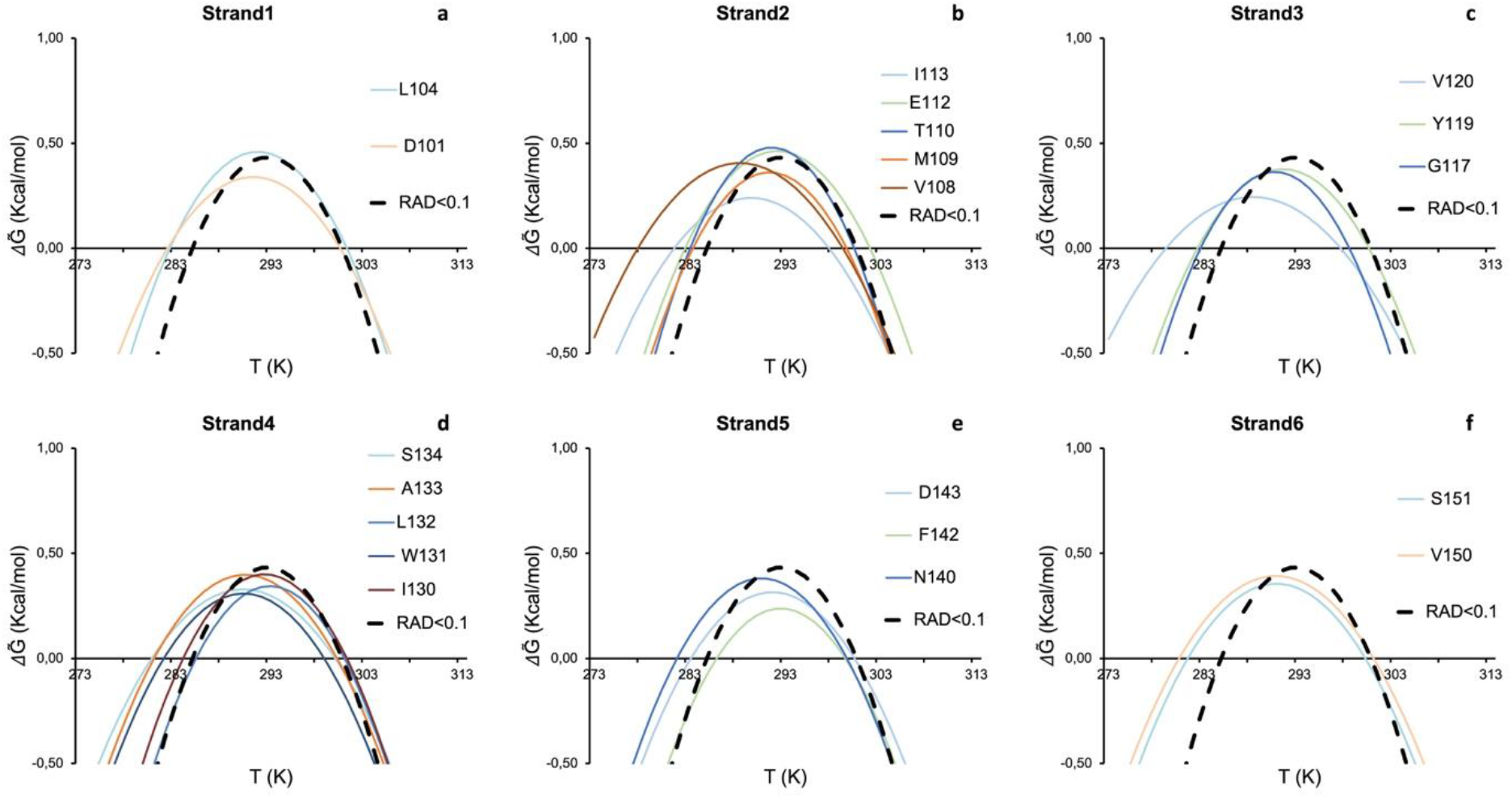
Stability curves of residues belonging to secondary structure elements. a) Strand 1. b) Strand 2. c) Strand 3. d) Strand 4. e) Strand 5. f) Strand 6. Residues are labelled with single letter code. The average stability curve is shown as black dashed line.

The best agreement was found for Leu104 of strand 1, Met109 and Thr110 of strand 2, Tyr119 for strand 3, Ile130 and Leu132 of strand 4 and Phe142 and Asp143 of strand 5.

### The behaviour of tryptophan side chains

We then looked into the possibility of following the process of unfolding and calculating thermodynamic parameters using the tryptophan side chains. This choice directly parallels studies based on following the process of unfolding by fluorescence using the intrinsic tryptophan fluorescence (Monsellier & Bedouelle, 2002). Yfh1 has two tryptophans: W131 is fully exposed to the solvent whereas W149 is buried. Both residues are fully conserved throughout the frataxin family and the two side chain resonances are clearly identifiable (**Figure S5a of Suppl. Mat.)**. We calculated the thermodynamic parameters for the side chain indole groups of both residues by the same procedure outlined for main chain NHs, generating first a stability curve. The resonance of W149, which could potentially be more interesting, could not be used for quantitative measurements because the temperature dependence of its volume yields a stability curve very different from the others (**Figure S5b of Suppl. Mat.)** and leads to impossible fitting parameters. This might be explained by the co-existence of folded and partially unfolded species in equilibrium with each other in solution. As a consequence, the indole of W149 resonates both at 9.25 and 127.00 ppm (folded species) and at ca. 10.05 and 129.20 ppm (split into three closely adjacent peaks, unfolding intermediates) (**Figure S5a of Suppl. Mat**.). As previously proven experimentally, the resonances of the unfolding intermediates disappear upon addition of salt (Figure 1, panel A and B in Vilanova et al., 2014). These resonances are also at the same chemical shifts observed for the tryptophan indole groups at low and high temperature where however the three signals collapse into one (**Figure S1 of Suppl. Mat**.). The complex equilibrium between different species could thus explain the ill-behaviour of the corresponding stability curve of this residue.

The behaviour of the resonance of the exposed W131 side chain is instead fully consistent with that of RAD_0.1 and also with the original curve calculated from 1D NMR data (Pastore et al., 2007) (**Table 1**). On the whole, these results exemplify well the complexity of the selection choice of the unfolding reporter and advocate in favour of a wholistic analysis of all the available data.

## Discussion

The *de facto* demonstration that it is possible to reliably measure the thermodynamic parameters of protein unfolding by 2D NMR spectroscopy (Puglisi et al., 2020) has opened a new territory to study protein unfolding at atomic resolution using site-specific information. Following protein folding/unfolding looking at specific residues rather than obtaining an average overall picture is not a novelty. Despite some intrinsic limitations, fluorescence has, for instance, been used for decades to probe protein unfolding following the intrinsic tryptophan fluorescence (Monsellier & Bedouelle, 2002; Bolis et al., 2004). Another elegant, although sadly still underexploited technique able to report local behaviour at the level of specific residues is chemically induced dynamic nuclear polarization (CIDNP), first introduced to the study of proteins by Robert Kaptein (Kaptein et al., 1978). This technique allows the selective observation of exposed tryptophans, histidines and tyrosines. In protein folding, it was, for instance, used to characterize the unfolded states of lysozyme (Broadhurst et al., 1991; Schlörb et al., 2006) and the molten globule folding intermediate of α-lactalbumin (Improta et al., 1995; Lyon et al., 2002). Real-time CIDNP was also used to study the refolding of ribonuclease A (Day et al., 2009) and HPr (Canet et al., 2003). The only drawback of this technique is that, as in fluorescence, the information is limited to specific aromatic residues.

Another important technique that reports on protein unfolding at the single residue level is stopped-flow methods coupled with NMR (Kim and Baldwin, 1991; Roder and Wüthrich, 1986) or mass spectrometry (Miranker et al., 1993) measurements of hydrogen exchange. In a classic paper (Miranker et al., 1991), Dobson and co-workers described, for instance, NMR experiments based on competition between hydrogen exchange as observed in COSY spectra and the refolding process. The authors concluded that the two structural domains of lysozyme followed two distinct folding pathways, which significantly differed in the extent of compactness in the early stages of folding. Similar and complementary conclusions could be reached by integrating NMR with mass spectrometry (Miranker et al., 1993). While these studies retain their solid importance, the possibility of following the resonance intensities also by HSQC spectra may provide a more flexible tool to obtain detailed information on unfolding, as this technique reports on the exchange regime but also, implicitly, on the chemical environment. The use of 2D HSQC had been discouraged by the non-linear relationship between peak intensity (or volume) and populations with temperature as the consequence of relaxation, imperfect pulses, and mismatch of the INEPT delay with specific J-couplings. We have previously suggested an approach to compensate for these effects and demonstrated that the non-linearity does not affect the spectra of Yfh1 (Puglisi et al., 2020), even though these conclusions might be protein dependent.

Here, we used the approach developed in our previous work (Puglisi et al., 2020) to analyse individual stability curves for most of the residues of Yfh1. Our analysis is highly complementary to the single residue information that may be obtained through HDX by NMR or mass spectrometry (Englander and Mayne, 1992; Miranker et al., 1996). A clear advantage of the current approach is the availability of signals of almost all residues and the relative simplicity of the analysis.

We noted that Yfh1 shows a multitude of events on top of the overall folding/unfolding. We observed that the behaviour of the individual stability curves is not distributed uniformly along the sequence. Residues can be clearly divided into two groups, i.e. those consistent with the average behaviour of an all-or-none mechanism of unfolding and those differing, even strongly, from the best average RAD_0.1. This finding alone proved that it is not possible to measure stability using a single residue without a careful evaluation of the role of the specific residue in the protein fold. This conclusion is partially mitigated by our results on the parameters obtained for a tryptophan indole. However, in the whole, also for these side chains it may be difficult, *a priori*, to infer which tryptophan is more reliable. We showed that, of the two tryptophans present in Yfh1 only the fully exposed W131 is suitable for the analysis. Our results thus demonstrate that unfolding studies based on fluorescent measurements using the intrinsic fluorescence of tryptophan should always be taken with a pinch of salt: in many cases no independent controls are feasible to evaluate the accuracy of the results. The possibility of using 2D NMR and the introduction of the easily approachable RAD parameter may assist in this choice in future studies.

Analysis of individual secondary structure elements, i.e. helices and strands, showed that there is no clear hierarchy among them, and there is no indication that any of the elements undergoes disruption before the others, either at high or low temperature. This implies that, overall, the folding/unfolding of the core of Yfh1 can be described as a single, highly cooperative event, but not all residues could be used for following the transition. It will be interesting in the future to study lysozyme to have an example in which two subdomains unfold independently (Miranker et al., 1991). In addition to information on regular secondary structure elements, our analysis yielded also interesting information on less ordered traits. Intrinsically flexible elements, i.e. regions characterized by multiple conformers, can be identified unequivocally by their thermodynamic parameters, without recurring to interpretative mechanisms.

Another important point is that we observed a clear difference between parameters corresponding to the cold and the heat denaturation processes: residues that are outliers from the average stability curve tend to have a strong stabilization effect at low temperature and a weaker destabilising effect at high temperature. This is a strong confirmation that the mechanisms of the two transitions are intrinsically different according to the mechanism of cold unfolding proposed by Privalov. In this model, cold denaturation is intimately linked to the hydration of hydrophobic residues of the core (Privalov, 1990) and with his suggestion that the disruption of the hydrophobic core at low temperature would be caused by the hydration of hydrophobic residue side chains of the core, whereas the high temperature transition is mainly linked to entropic factors, consistent with the increase of thermal motions when temperature is increased. This is what we observed in our NMR analysis of Yfh1 and is in line with our previous evidence that showed that the unfolded species at low temperature has a volume higher than the folded species and of the high temperature unfolded species (Alfano et al., 2017) and that cold denaturation is caused by a hydration increase (Adrover et al., 2012).

We also observed, more surprising, that some residues not belonging to the hydrophobic core have T_c_s appreciably lower than the average. A possible explanation for this behaviour is that, at the temperature of global unfolding, corresponding to that of the average RAD_0.1 of the deeply buried protein core, residues outside the hydrophobic core and in regions classified as flexible could be more resilient against unfolding. This would imply that, at low temperature, opening of the hydrophobic core and its disruption could happen before the collapse of external and more exposed elements: the core would unfold in lowering the temperature whereas outer turns could be affected last.

## Conclusions

In conclusion, we have provided here a nice example of a protein that only apparently follows a simple two-state (folded/unfolded) statistical model and for which a global denaturation curve simply hides a profound intrinsic heterogeneity. We described in detail how the unfolding of Yfh1 is a much more complex process than a two-step global unfolding both at high and low temperature. Our data clearly show how, as recently advocated by Grassein et al. (2020), local nativeness is probe dependent and, as such, needs to be studied at the individual residue level. The possibility of studying the process relied in our case on the nearly unique properties of Yfh1 but also, more in general, on the use of NMR which is probably the most suitable technique to analyse the contributions to the (un)folding process in a residue-specific manner. We can certainly state that monitoring protein unfolding by the stability curves of individual residues, as allowed by 2D NMR spectroscopy, yielded a much more informative picture than what may have been obtained by any other traditional method. Our work thus paves a new way to the study of protein unfolding that will need to be explored in the future using a number of completely different systems to reconstruct a more complete picture of the complexity of the process.

## Experimental session

### Sample preparation

Yeast frataxin (Yfh1) was expressed in BL21(DE3) *E. coli* as previously described (Pastore et al., 2007). To obtain uniformly ^15^N-enriched Yfh1, bacteria were grown in M9 using ^15^N-ammonium sulphate as the only source of nitrogen until an OD of 0.6-0.8 was reached and induced for 4 hours at 310K with IPTG. Purification required two precipitation steps with ammonium sulphate and dialysis followed by anion exchange chromatography using a Q-sepharose column with a NaCl gradient. After dialysis the protein was further purified by a chromatography using a Phenyl Sepharose column with a decreasing gradient of ammonium sulphate.

### NMR measurements

2D NMR ^15^N-HSQC experiments were run on a 700 MHz Bruker AVANCE spectrometer. ^15^N-labelled Yfh1 was dissolved in 10 mM Hepes at pH 7.5 to reach 0.1 mM with 0.1 mM selectively 15N-labelled tyrosine CyaY. Spectra were recorded in the range 278-313 K with intervals of 2.5 K and using the Watergate water suppression sequence (Piotto et al., 1992). For each increment 8 scans were accumulated, for a total of 240 increments (TD). Spectra were processed with NMRPipe and analysed with CCPNMR software. Gaussian (LB -15 and GB 0.1) and cosine window functions were applied for the direct and indirect dimension respectively. The data were zero-filled twice in both dimensions. Spectral assignments of Yfh1 were taken from the BMRB deposition entry 19991.

### Selection of the amides to be used in our analysis

Yfh1 contains 114 backbone amide protons. The first 23 residues are intrinsically disordered (Popovic et al., 2015) and are part of the signal peptide for mitochondrial import, leading to 91 resonances in the globular domain. Sixty eight residues have non-overlapping and isolated resonances that allow easily detectable and reliable volume calculation. Most of the excluded overlapping resonances corresponded to disordered regions or to partially unfolded conformations in equilibrium with the folded one in a slow exchange regime at room temperature (Sanfelice et al., 2014).

### Calculations of the RAD parameters

The RAD parameter of the backbone amide nitrogen atoms of Yfh1 was calculated on the crystallographic coordinates of a Tyr73-to-Ala mutant solved at 3.0 A resolution (2fql, Kalberg et al., 2006). This choice was dictated by the better resolution of this structure as compared to an alternative NMR structure (2ga5) or to homology models. The mutation, that is at the very beginning of the globular region of the protein, does not affect the structure of the protein as demonstrated by comparison with other orthologs but changes the self-assembly properties of the protein (Kalberg et al., 2006). No hydrogen atoms were added. RAD was obtained using the software Pops (https://github.com/mathbio-nimr-mrc-ac-uk/POPS) and SADIC (http://www.sbl.unisi.it/prococoa/). As previously described (Puglisi et al., 2020), the RAD parameter was defined according to the equation

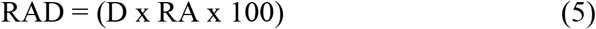

where D was the distance of an atom from the protein surface as calculated by the program SADIC (Varrazzo et al., 2005). RA was the relative accessibility at atomic level RA defined as the ratio between the exposed surface of a nitrogen atom with respect to that of the whole residue and calculated by the software POP (Cavallo et al., 2003). Most of the exposed residues had RAD values for the amide nitrogens considerably higher than 0.5 and were excluded from the analysis (**Table 1**). The curves obtained for individual resonances using RAD values between 0.5 and 0.1 had a lower relative spread and a much better agreement with the CD curve (**data not shown**). The stability curve and the thermodynamics parameters calculated from averaging amide volumes from residues with a RAD value below 0.1 (RAD_0.1) were fully consistent with those calculated from CD spectroscopy, within experimental error (Puglisi et al., 2020). Residues involved in secondary structures were evaluated according to the DSSP program (https://swift.cmbi.umcn.nl/gv/dssp/).

### Calculation of the stability curves

Volumes were calculated by summation of the intensities in a set box using the CCPNMR software (https://www.ccpn.ac.uk/v2-software/software). The volumes were normalized by dividing the volume of each peak of Yfh1 at a given temperature by the volume of CyaY Tyr69 amide peak at the same temperature as previously described (Puglisi et al., 2020). This normalization is meant to filter out the non-linearity of the relationship between peak intensity (or volume) and populations due to instrumental effects. The corrected volumes were transformed into relative populations of folded Yfh1.

At each temperature, the fraction of folded protein was estimated by the equation

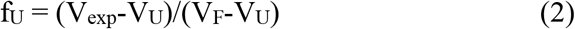

where V_exp_ is the measured volume, V_U_ is the volume of the unfolded state (assumed at 313 K), and V_F_ is the volume of the folded (maximum value) taking into account that, as previously proven (Pastore et al., 2007), at room temperature the unfolded forms of Yfh1 are in equilibrium with the folded population present on average at 70%.

The fraction of folded, f_F_(T), and unfolded, f_U_(T), forms are a function of the Gibbs free energy of unfolding, ΔG°(T). If the heat capacity difference between the folded and unfolded forms, ΔC_p_, is assumed independent of temperature, the free energy is given by the Gibbs-Helmholtz equation (Martin et al., 2008). The thermodynamic parameters T_m_, ΔH_m_ and ΔC_p_ were derived by nonlinear least-squares fitting using the Levenberg-Marquardt algorithm from the following equation and omitting the points at 313 K for which, by definition from our assumption, f_U_ is equal to 1.

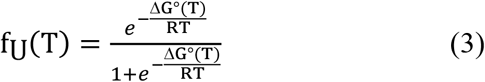

in which T_m_, ΔH_m_ and ΔC_p_ can be obtained by fitting the modified Gibbs-Helmholtz equation

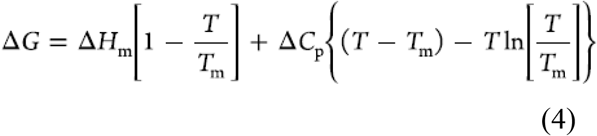

The curve corresponding to this equation is known as the stability curve of the protein (Becktel and Schellman, 1987). Other parameters for low temperature unfolding, e.g. the low temperature unfolding (T_c_), were obtained from the stability curve.

Errors on the stability curves were evaluated propagating the errors from the covariance matrix of the fit. In the representative fits reported in **Suppl. Mat**. (**Figures S6-S8**), errors were represented as gray lines calculated by the covariance method (Press et al., 1988). They represent how well the measured populations (and thus ΔG) vs. temperature agree with the equation for the stability curve. We reported six representative curves from the subset used to calculate RAD_0.1 (**Figure S6**), four curves from the subset of Figure 1c (**Figure S7**), and four curves corresponding to the best-behaved residues of the beta sheet (**Figure S8**). The curves do not fully represent ΔG because, despite we assumed the protein completely unfolded at 313K, fitting showed that not all the residues reached a plateau of unfolding at high temperature. We thus indicated the curves as 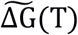 to underline the distinction.

## Acknowledgments

This manuscript is meant in celebration of the 80^th^ birthday of Prof. Robert Kaptein. We thank Geoff Kelly and Tom Frenkiel of the MRC Biomedical NMR Centre for helpful discussions and technical support, Neri Niccolai and Franca Fraternali for help with their software SADIC and PopS respectively. We also thankfully acknowledge the use of the NMR spectrometers at the Randall unit of King’s College London and the Francis Crick Institute for access to the MRC Biomedical NMR Centre. The Crick Institute receives its core funding from Cancer Research UK (FC001029), the UK Medical Research Council (FC001029) and the Wellcome Trust (FC001029). The research was supported by UK Dementia Research Institute (RE1 3556) that is funded by the Medical Research Council, Alzheimer’s Society and Alzheimer’s Research UK.

## Notes

### Competing Interest Statement

The authors have declared no competing interest.

